# Time-resolved digital immunoassay for rapid and sensitive quantitation of procalcitonin with plasmonic imaging

**DOI:** 10.1101/483347

**Authors:** Wenwen Jing, Yan Wang, Yunze Yang, Yi Wang, Guangzhong Ma, Shaopeng Wang, Nongjian Tao

**Author notes:** These authors contributed equally to this work (Wenwen Jing, Yan Wang).

## Abstract

Timely diagnosis of acute diseases improves treatment outcomes and saves lives, but it requires fast and precision quantification of biomarkers. Here we report a time-resolved digital immunoassay based on plasmonic imaging of binding of single nanoparticles to biomarkers captured on a sensor surface. The real-time and high contrast of plasmonic imaging lead to fast and precise counting of the individual biomarkers over a wide dynamic range. We demonstrated the detection principle, evaluated the performance of the method using procalcitonin (PCT) as an example, and achieved a limit of detection of ~ 3 pg/mL, dynamic range of 4-12500 pg/mL, for a total detection time of ~ 25 mins.

## Introduction

Detection and quantification of molecular biomarkers are critical to disease diagnosis and progression monitoring. ^[1-3]^ Various approaches have been developed ^[4-7]^, but the most well-established technology is ELISA (enzyme-linked immunosorbent assay), which amplifies antibody-biomarker binding via enzymatic reactions and converts reaction products into an optical signal (e.g., color changes) ^[8-9]^Although ELISA is widely used in clinical and research labs, its detection limit, total test time (sample to result) and dynamic range are often insufficient for clinical applications. Recent technological advances have made it possible to detect single molecules, which have been used to measure the binding of a single biomarker molecule to a capture antibody with improved detection limit. This single-molecule approach has been referred to as digital immunoassay ^[10-13]^

Several digital immunoassay methods have been developed or proposed. One is to replace the large reaction well in the traditional ELISA with an array of microwells ^[10-13]^The volume of the microwells is small, such that each microwell has either no biomarker present or a single biomarker molecule that binds to a capture antibody conjugated with a magnetic bead in the microwell. Another digital immunoassay platform combines single molecule fluorescent detection with flow cytometry [14]. The third platform uses metal nanoparticles as the signal readout mechanism ^[15-18]^. The nanoparticles can be imaged individually via dark field, interference and plasmonic optical imaging techniques ^[19-20]^These digital immunoassay technologies provide single molecule detection capability, which significantly improves the detection limit of the traditional ELISA. However, the improvement is often achieved at the expense of test time (e.g., > 1 hr) and dynamic range (e.g., < 2 ng/mL), both are crucial for disease diagnosis and treatment ^[21]^An important example is sepsis, which develops rapidly with a survival rate of 79.9% from the first hour and a sharp drop of 7.6% per hour ^[22]^Timely detection of biomarkers of sepsis is thus needed to improve treatment outcome and save lives.

Here we report a time-resolved digital immunoassay (TD-Immunoassay) based on plasmonic imaging of nanoparticles for rapid detection of biomarkers with a wide dynamic range. The plasmonic imaging offers high contrast and fast imaging of nanoparticles, allowing detection of single molecule binding on a sensor surface via detection of antibody-conjugated nanoparticles. It features real-time counting of the nanoparticles as they bind to the biomarker molecules captured by antibody, which provides accurate assessment of the biomarker concentration without the need of reaching thermal equilibrium via lengthy incubation. The real-time counting together with super-localization tracking of each nanoparticle allows resolving two binding events within a distance smaller than the diffraction limit ^[23]^, which enhances the dynamic range and minimizing the counting error. Using TD-Immunoassay, we have achieved a limit of detection of ~ 3 pg/mL, dynamic range of 4-12500 pg/mL, and total detection time of ~ 25 mins for procalcitonin (PCT), an important biomarker for sepsis ^[24]^

## Materials and methods

### Chemicals

N-hydroxysuccinimide (NHS), N-ethyl-N’-(3-dimethylaminopropyl) carbodiimide hydrochloride (EDC), sodium acetate (NaOAc), acetic acid (AcOH), Tween 20, rabbit Immunoglobulin G (IgG) and bovine serum albumin (BSA) were purchased from Sigma-Aldrich (St. Louis, MO). 150 nm gold nanoparticles coated with goat anti-rabbit IgG (C11-150-TGARG-50), and 150 nm gold nanoparticles coated with streptavidin (C11-150-TS-DIH-50) were from Nanopartz™. Dithiolalkanearomatic-PEG3-OH (Dithiol-PEG-OH) and dithiolalkanearomatic-PEG6-COOH (Dithiol-PEG-COOH) were purchased from SensoPath Technologies (Bozeman, MT). Human PCT ELISA kit (CATALOG NUMBER: dy8350-05), and DuoSet^®^ ancillary reagent kit 2 (catalog number: DY008) were purchased from R&D system, USA. Human serum (from human male AB plasma, USA origin, sterile-filtered) was purchased from Sigma-Aldrich, USA. The kits and reagents mentioned above were sub-packed and stored according to the instructions.

### Plasmonic imaging

Plasmonic imaging was implemented on an inverted optical microscope (IX- 81, Olympus, Shinjuku, Tokyo, Japan) with a 60x high numerical aperture (NA 1.49) oil immersion objective. A collimated p-polarized light beam (1 mW) from a 680 nm fiber coupled super luminescence diode (from Qphotonics, Ann Arbor, MI) was directed onto a sensor surface via the objective to excite surface plasmons. The plasmonic images were collected by a CCD camera (Pike, F-032B, Allied Vision Technologies, Newburyport, MA) at a frame of 106 frames per second with 4 frames binning (effective frame rate, 26.6 fps) with a view area of 640 × 480 pixels and a pixel size of 7.4 μm. The sensors were prepared by coating BK-7 glass coverslips with 1.5-2 nm chromium, followed by 47 nm gold. A Flexi-Perm silicone solution cell (SARSTEDT, Germany) was placed on top of the sensor to hold the sample solution. Automated particle counting from the images were performed with an imaging processing algorithm described in the supporting information (section S2).

### Sample preparation

50 μg/mL rabbit IgG solution was prepared by dissolving rabbit IgG in NaOAc/AcOH buffer (10 mM pH=5.0 NaOAc/AcOH). The solution was diluted via multiple serial dilution, 10x each, to reach different concentrations of rabbit IgG, from 5×10^-9^ to 5×10^-1^μg/mL. PCT capture antibody (240 μg/mL) stock solution was diluted with NaOAc/AcOH buffer (10 mM pH=5.0 NaOAc/AcOH) to reach 2 μg/mL. PCT standard was diluted with reagent diluent (solution from DuoSet^®^ ancillary reagent kit 2) using double ratio dilution method, to reach final of 12500, 6250, 2000, 1000, 200, 31.3, 7.83, 3.9, and 1.95 pg/mL. PCT spiked serum samples were prepared by diluting the PCT standard into human serum to reach 2000, 200, and 20 pg/mL. PCT detection antibody (3 μg/mL) stock solution was diluted with reagent diluent to reach 50.00 ng/mL. 150 nm goat anti-rabbit IgG coated gold nanoparticle solution was prepared by diluting the stock solution 1000 times with DI water and then sonicated for 5 min. Streptavidin coated gold nanoparticle solution was prepared by adding 15 μL of gold nanoparticle solution to 135 μL of PBS buffer and then sonicated for 5 min.

### Sensor surface modification

The sensors were cleaned with deionized water (Milli-Q, Millipore Corp.) and then ethanol, followed by hydrogen flaming. After cleaning, the sensors were immediately soaked into 1 mM of 50:1 PEG-OH/PEG-COOH mixed dithiol ethanol solution for 24 hours in dark. The sensors were then rinsed with deionized water and ethanol, and then dried with nitrogen gas. Each sensor was activated for immobilization of a receptor molecule to its surface by adding 100 μL NHS/EDC aqueous solution (containing 100 mM NHS and 400 mM EDC), followed by gradient washing (gradually diluted the NHS/EDC solution by DI water).

### IgG binding to anti-IgG

IgG was immobilized to an activated sensor surface by incubating the sensor in 100 μL of prepared rabbit IgG solution at room temperature for 10 min, followed by washing it with 250 μL of PBST buffer (0.05% Tween 20 in PBS buffer, Corning Cellgro) three times. Residual activated binding sites were blocked by 250 μL BSA solution (1% w/v) for 5 min and then washed with 250 μL of PBST buffer three times. 100 μL solution of gold nanoparticles coated with goat anti-rabbit IgG was added, and the binding of the gold nanoparticles to IgG was tracked in real time.

### Detection of PCT

PCT capture antibody was immobilized to an activated sensor by incubation with 100 μL PCT capture antibody solution for 10 min at room temperature, followed by washing with 250 μL of PBST three times. Residual activated binding sites were blocked by 250 μL of BSA solution (3% w/v) for 5 min and then washed with 250 μL of PBST buffer three times. The capture antibody-functionalized sensor was either used immediately or stored at 42°C prior to PCT assay. The detailed protocol of PCT immune-sandwich assay was described in the Supporting Information (Fig. S1).

### Conventional ELISA assay for PCT

Following the manufacturer’s protocol, 100 μL capture antibody was added to each well of a microplate and incubated overnight at room temperature. Next, the microplate was washed with PBST buffer for three times, and then blocked with 300μ L reagent diluent for at least 1 hour. After washing three times with PBST buffer, 100 μL standard or sample was added and incubated for 2 hr at room temperature, and then washed three times. Next, 100 μL detection antibody was added and incubated for 2 hr, and then washed three times. Next, 100 μL of streptavidin-HRP was added and incubated for 20 min followed by three times washing. Next, 100 μL substrate solution was added to the well and incubated for 20 min, and finally 50 μL stop solution was added. The optical density at 450 nm was measured using a microplate reader (model: Varioskan Flash, ThermoFisher Scientific).

### Statistical Analysis

We use linear regression model for PCT counting error prediction and 95% prediction interval to evaluate the prediction accuracy from the counting numbers (see supporting information for details). Data is processed by custom-written Matlab scripts.

## Results

### Principle and setup

A gold-coated glass slide was used as the sensor, on which a capture antibody for PCT (procalcitonin) was immobilized via a PEG linker (Fig. 1a). A sample was introduced to allow binding of PCT in the sample to the capture antibody. A detection antibody, biotinylated anti-PCT, was then added to bind with the PCT bound to the capture antibody and form a capture antibody- PCT-detection antibody complex, followed by introducing streptavidin-coated gold nanoparticles (GNP) with a diameter of 150 nm, which bound to the antibody complex via the biotin-streptavidin interaction. The surface density of the capture antibody was adjusted such that each gold nanoparticle could bind only with one PCT molecule, providing a readout of a PCT binding event on the surface.

**Figure 1:**
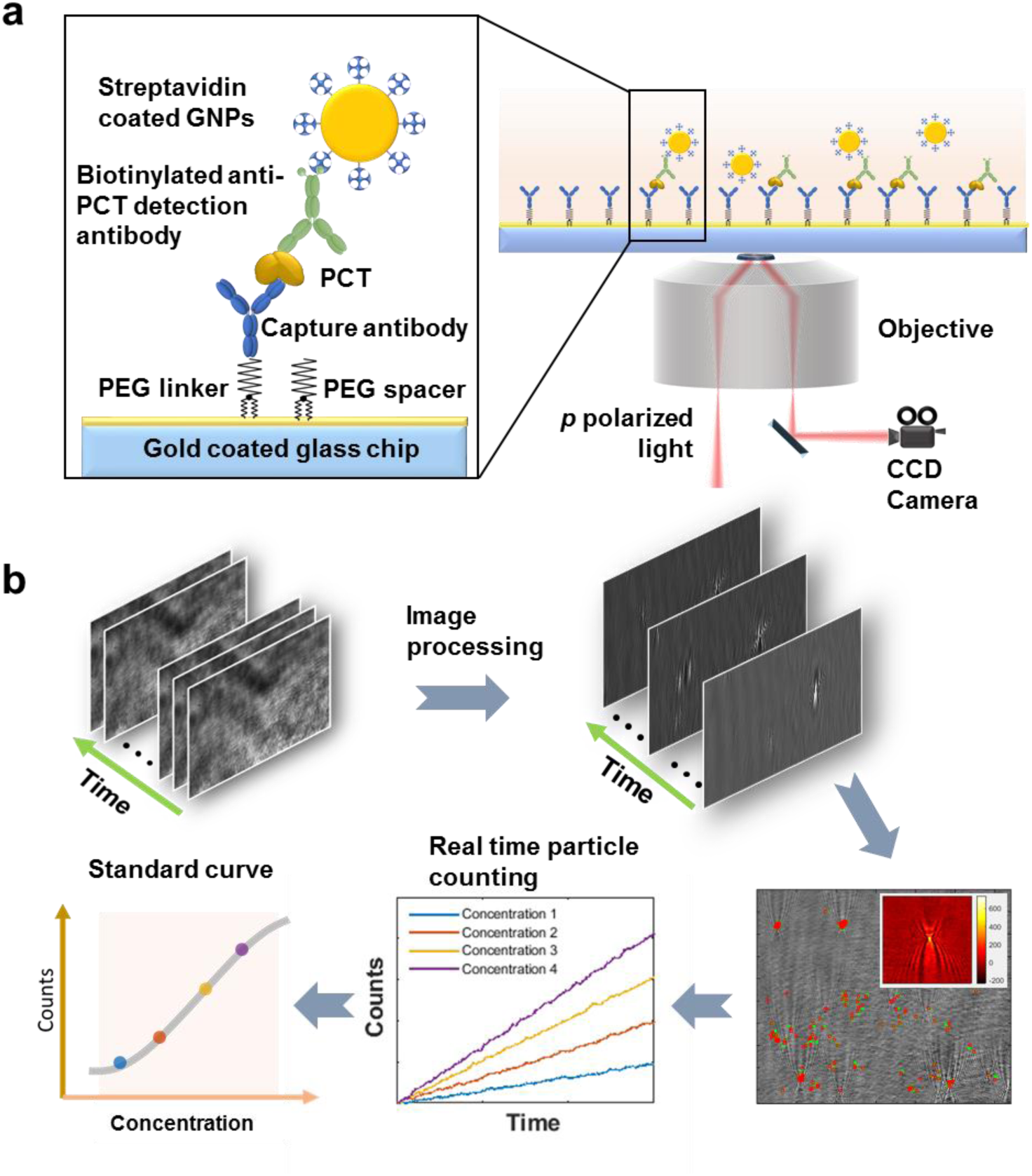
Schematic illustration of time-resolved digital immunoassay. **(a)** Experimental setup. A SPR sensor chip (gold-coated glass slide) is functionalized with capture antibody via polyethylene glycol (PEG) linker. PCT in the sample solution binds to the capture antibody, which is detected by gold nanoparticles that bind to a detection antibody that binds to the PCT on the capture antibody. The binding processes are monitored with a plasmonic imaging system built on an inverted optical microscope with details described in the text. **(b)** Workflow of imaging and data processing, which removes background noise from the raw plasmonic images via differential imaging, tracks binding and binding of the gold nanoparticles to the sensor surface, determines the net count of bound gold nanoparticles vs. time, and generate a standard curve (binding events. Vs. PCT concentration).

Sensitive detection of single GNPs was achieved with a plasmonic imaging platform, where a p polarized light beam with a proper incident angle from a superluminescence (SLED) diode was directed onto the sensor surface via a 60x high numerical aperture oil immersion objective to excite plasmons on the gold surface. The scattered and reflected light was collected with the same objective and imaged with a CCD camera. The time sequence of the plasmonic images captured the binding of the individual GNPs with a temporal resolution of ~ 37.5 ms (Fig. 1b). Each GNP was revealed as a bright spot with a parabolic pattern arising from the scattering for the plasmonic wave on the sensor surface by the GNP (colored inset at lower right of Fig. 1), which provides high image contrast and facilitates accurate tracking of single GNPs ^[25^,^26]^. Using an automated imaging processing algorithm, we tracked the position of each individual GNPs and counted the individual GNPs binding to the surface over time, from which a standard curve of PCT was obtained for concentration calculation (Fig. 1b).

The automated particle counting algorithm consists of following steps. First, noise in the image sequence is reduced by performing spatial Fast Fourier transform, and then differential images are obtained by subtracting the first image frame from the rest of the frames, followed by time average over multiple frames ^[27]^. These procedures remove common noise in the optical system and provides high contrast images of single GNPs (Fig. 2a). Second, the algorithm uses the distinct parabolic pattern of a single GNP plasmonic image as a template to search and identify all the GNPs on the sensor surface with an autocorrelation pattern recognition algorithm (see supporting information for more details). Because both the binding and unbinding of the GNPs take place on the sensor surface dynamically, two opposite patterns are observed for the binding (top of Fig. 2b) and unbinding events (bottom of Fig. 2b), respectively. The former is: GNP image subtract background, and the latter is: background subtract GNP image, so the images of binding and unbinding events are inverted in contrast. This procedure allows the algorithm to differentiate and track both binding and unbinding processes over time (Fig 2c). Third, the spatial location of each GNP was determined and tracked with a procedure described in the Supporting information (Fig. S2). Super-resolution fluorescent microscopy was used by Gooding et al. to track binding events within a distance smaller than the optical diffraction limit. ^[20]^ This is achieved with the dynamic tracking capability of the plasmonic imaging in the present work, which resolves multiple binding events within an area smaller than the optical diffraction limit in time domain, as long as no two or more GNPs bind to the area at the same moment (defined by the frame rate). This capability is critical for improving the counting accuracy and expanding the dynamic range.

**Figure 2:**
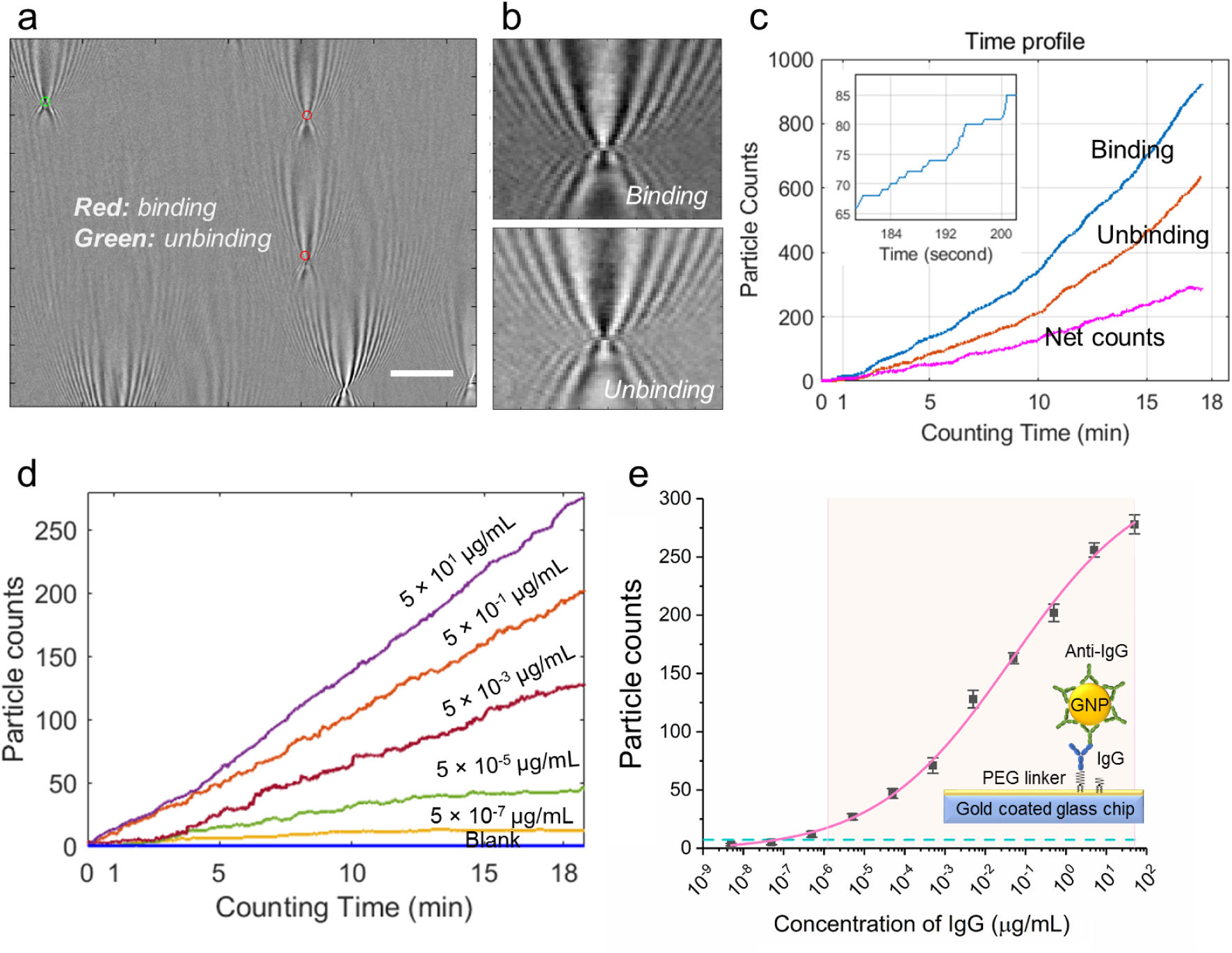
IgG/Anti-IgG binding quantification. (a) A typical differential plasmonic image, showing both binding (red) and unbinding (green) of gold nanoparticles (diameter, 150 nm). Scale bar: 10 μm (b). Zooming-in of binding and unbinding gold nanoparticles, showing inverted contrast in the differential images. (c) Binding (blue), unbinding and net (magenta) counts of gold nanoparticles vs. time with Anti-IgG-gold nanoparticle concentration of 50 μg/mL. Figure in the upper left shows a zoom in zigzag plot of binding curve. (d) Nanoparticle counts (net) vs. incubation time at different IgG concentrations. For clarity, only data of some concentrations are plotted here. (e) Standard curve of IgG detection, where the error bars are the standard deviation from triplicate tests, the curve is a fit to the equation shown in the supporting information, the dashed horizontal line is GNP counts of blank solutions, and the shaded area mark the dynamic range.

### Validation of TD-immunoassay with IgG/anti-IgG binding

To validate the capability of TD-immunoassay for detecting antigen and quantifying antigen concentration, we first applied it to study the binding of IgG binding to anti-IgG. IgG was immobilized onto the sensor surface, followed by incubation with anti-IgG conjugated GNPs at different concentrations and the binding of the anti-IgG to the IgG on the sensor surface was tracked by counting the individual GNP binding events on the surface with a temporal resolution of ~ 37.5 ms. The number of GNPs counted with the algorithm varies with concentration, but for each concentration it increases linearly with time (Fig. 2d). This indicates that binding/unbinding process within the time interval is far from reaching saturation of the binding sites (IgG) on the sensor surface by the anti-IgG, and also far from reaching thermal equilibrium.

From the GNP counts obtained at 20 min, we obtained a standard curve of IgG detection (Fig. 2e). We repeated the experiment 3 times and found less than 20% variability for each concentration. From the standard deviation of the triplicate tests, error bars were determined and marked in Fig. 2e. The limit of detection was determined by the mean concentration measured for the blank solutions plus three times of standard deviations ^[28]^, which is 64.6 fg/mL (or 0.43 fM, blue dashed line of Fig. 2e). The detectable dynamic range was from 1.3 pg/mL to 50 μg/mL, covering 7 orders of magnitude. These results demonstrate the feasibility and performance of TD-immunoassay.

### PCT detection

Detection of PCT was based on sandwich assay, where a sample containing PCT was introduced and incubated for 10 min to allow binding of PCT to the capture antibody immobilized on the sensor surface (see supporting information for detailed procedure). Biotinylated detection antibody was then introduced followed by introducing streptavidin-GNPs, which bind to the detection antibody via the biotin-streptavidin interaction. The binding process was tracked over time to determine the PCT concentration in each sample. Fig. 3a shows the GNP counts for different concentrations of PCT in reagent diluent buffer over 20 min recorded with a frame rate of 26.6 fps. The PCT concentration ranges from 0 (blank) to 1.25×10^4^ pg/mL, which covers the concentration range of sepsis patients (0-10^4^ pg/mL). The dynamic range of the present TD- immunoassay estimated from the maximum density of GNPs is significantly higher than demonstrated here (see discussion). Fig. 3b shows the standard curve of the PCT test at the 20- minute time point, showing a linear response of 4 logs ranging from 4 pg/mL to 1.25×10^4^ pg/mL (r-square = 0.9981). The error bars in the standard curve are the standard deviations over triplicate tests. The limit of detection determined by the mean concentration measured for the blank solutions plus three times of standard deviation is 2.76 pg/mL, which is marked by the horizontal dashed red line in Fig. 3b. The limit of quantification defined by the mean concentration measured for the blank solutions plus ten times of the standard deviation is 4 pg/mL. This performance is sufficient to cover sepsis detection, where the PCT level varies from 50 to 10000 pg/mL.

**Figure 3:**
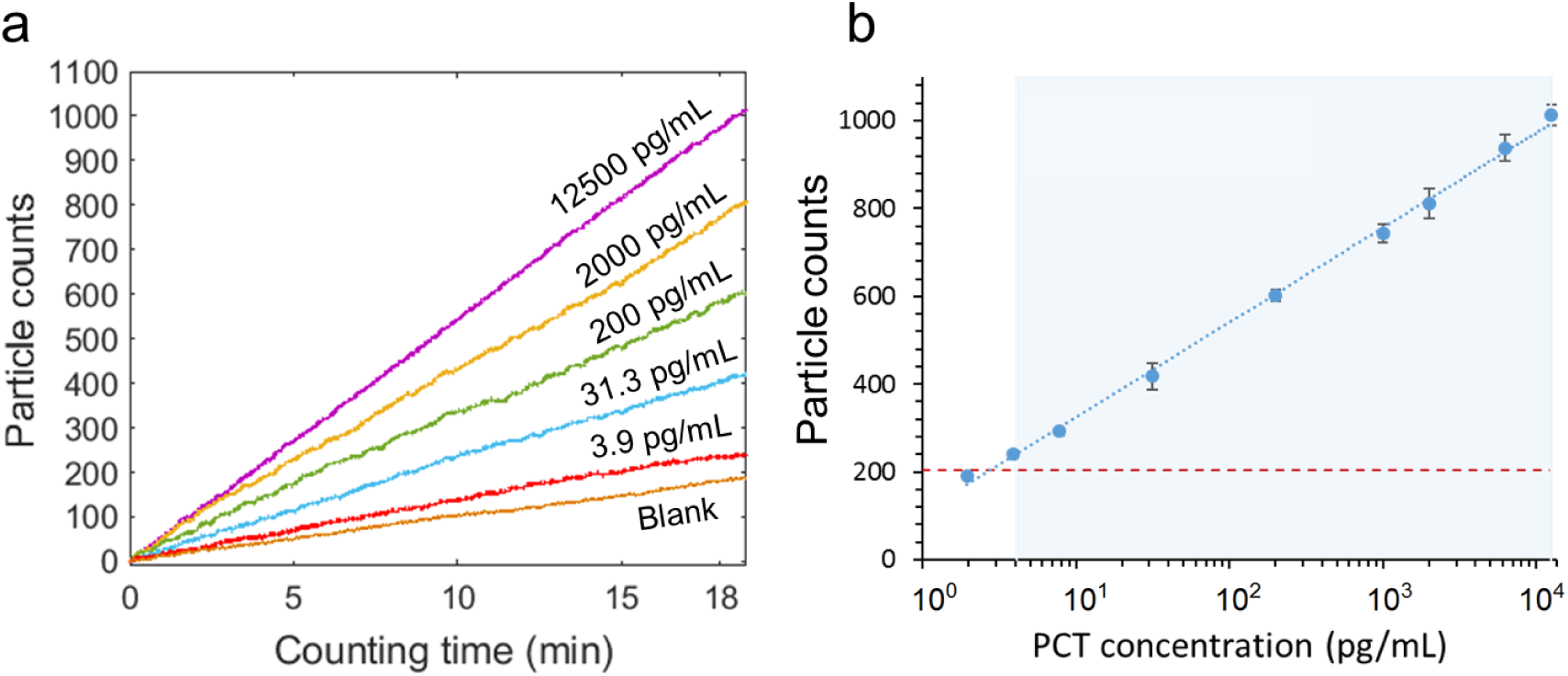
PCT detection. (a) Gold nanoparticle counts vs. binding time at different PCT concentrations. For clarity, only data of some concentrations are plotted here. (b) Standard curve of PCT detection, where the error bars are the standard deviation from triplicate tests, the curve is a fit to the equation shown in the supporting information, the dashed horizontal line is GNP counts of blank solutions, and the shaded area mark the dynamic range.

For comparison, the standard curve (Fig. S4) obtained by the conventional ELISA for PCT shows a logarithmic response with a dynamic range of 62.6 pg/mL to 2000 pg/mL and a limit of detection of 31.3 pg/mL. The standard curve from the present plasmonic imaging-based TD- immunoassay has a broader dynamic range (4 logs vs. 2 logs), a lower limit of detection (2.76 pg/mL vs. 31.3 pg/mL) and a lower limit of quantitation (4 pg/mL vs. 62.6 pg/mL).

The time-resolved detection of single GNPs helps improve the dynamic range and minimizing detection error associated with the binding of multiple GNPs to an area smaller than the spatial resolution of optical imaging. We show below that the real-time detection also helps to optimize the detection time required to achieve a desirable detection limit and precision. We studied the detection limit and precision by measuring GNP count vs. PCT concentration with GNP counting time of 1, 2, 5 and 15 min, respectively (Fig. 4). Three replicates were carried out for each concentration and for each GNP counting time, which are shown as black dots in Fig. 4. As expected, shorter GNP counting time leads to more scattered data due to smaller number of GNPs counted, but the GNP count is proportional to the logarithm of PCT concentration in each case. To evaluate the detection limit and concentration precision, we used a statistical model (see materials and methods, statistical analysis) to determine 95% prediction intervals (marked by the blue and red shaded regions in Fig. 4). The analysis shows that the error decreases with increasing GNP counting time, but the 5-min error is about the same as the 15-min error. This indicates that 5-min GNP counting is sufficient for precise PCT quantification.

**Figure 4:**
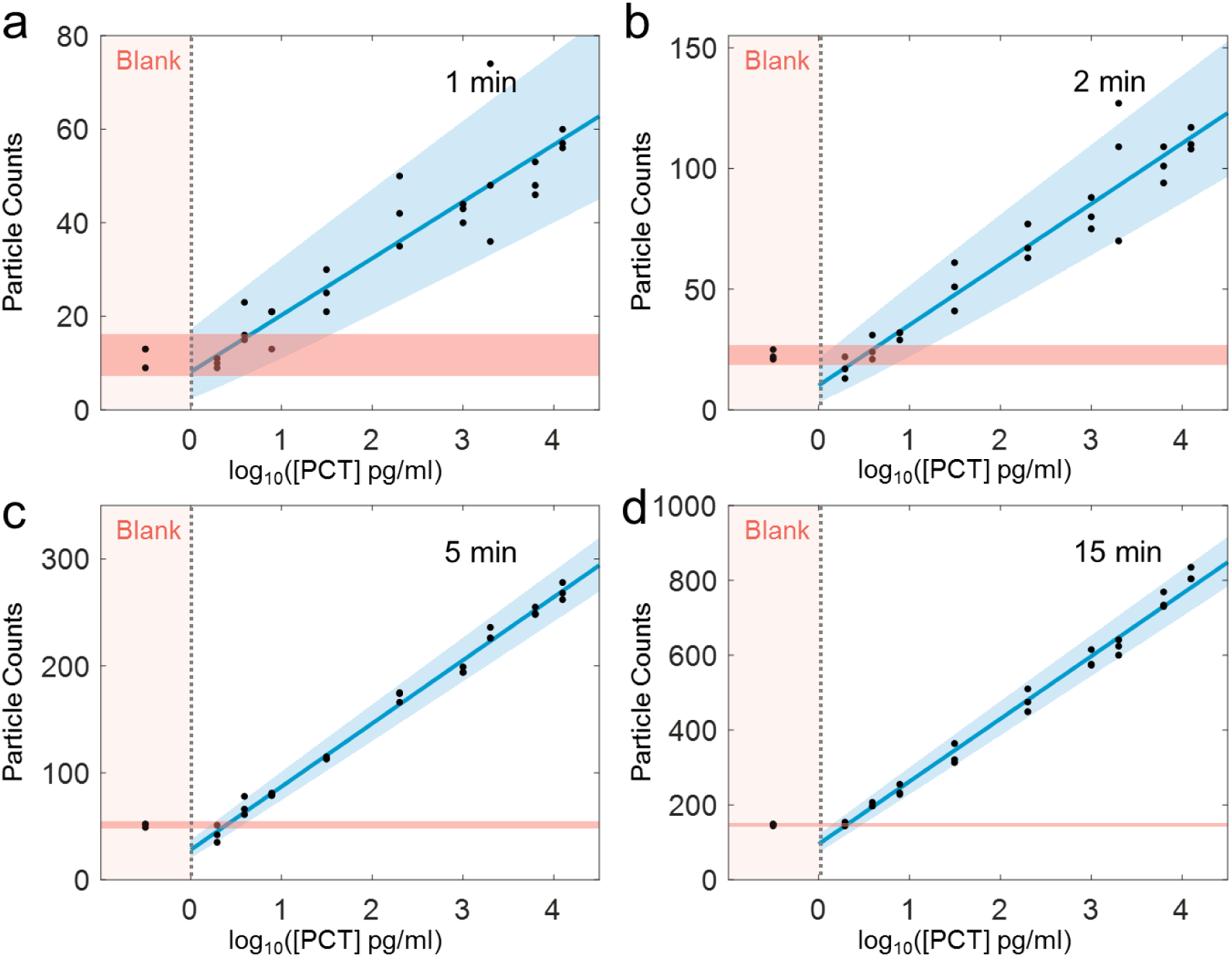
Particle counts vs. PCT concentration at different counting time intervals. TD- immunoassay measurements of PCT at different concentrations and comparison with blank solutions for 1 (a), 2 (b), 5 (c), 15 (d) min of gold nanoparticle counting time intervals. The counts of measurement are proportional to logarithm of PCT concentration, where the solid blue line is linear fitting to the data, the light blue and light red regions indicate 95% prediction interval according to the statistic model (materials and methods, statistical analysis) and measurement data. Three replicates for each concentration was carried out.

We further examined the detection limit and precision of TD-immunoassay by comparing the GNP counts over time at different PCT concentrations, including blank solutions (Figs. 5a-d). The data shows once again that the detection limit and precision improve with GNP counting time. The data also confirms that 5 min detection time can lead to detection of PCT concentration of 3.9 pg/mL. We evaluated the limit of quantification and prediction accuracy vs. time, which shows high precision data can be obtained within 5 mins, and longer time does not significantly improve the precision (Figs. 5e-f). Considering that the experiment involved 10 min incubation for PCT binding to the capture antibody and 10 min incubation for detection antibody binding to the PCT bound to the capture antibody, the total detection time is 25 min. The detection time achieved here compares favorably with other high-performance detection techniques (Table 2) and may be further shortened by eliminating the second incubation step using detection-antibody conjugated GNPs. It is worth mentioning that the higher concentration of the biomarker the shorter the overall detection time will be. This provides a possibility of detecting a biomarker using only the time required to reach a fixed precision, rather than using a fixed time interval regardless the biomarker concentration.

**Table 2.** Comparison of different PCT detection technologies

|  | TD-immunoassay<br>(This work) | Convention<br>al ELISA | ELECSYS<br>BRAHMS PCT<br>(Roche) | Quanterix<br>SiMoA <sup>[29]</sup> | SMCxPRO™<br>(MilliporeSigma) <sup>1</sup> |
| --- | --- | --- | --- | --- | --- |
| Sample volume (μL) | 100 | 100 | 30 | 100 | 1.6-3 |
| Incubation time | 10 min | 2 hour | 18 min | 20 min | 1 h |
| Total assay time | 25 min | 1-2 days | 3 h | > 3 h | 1 day |
| LoD (pg/ml) | 2.8 | 30 | 20 | 0.44 | Not available |
| LoQ (pg/ml) | 4 | >30 | 60 | 1.23 | 0.05-2 |
| Dynamic range (pg/ml) | 4-12500 | 31.3-2000 | 20-10000 | 1.23-900 | > 4 logs |

<sup>1</sup>The parameters of SMCxPRO™ was the theoretical value of the instrument, not specific obtained from PCT detection.

**Figure 5:**
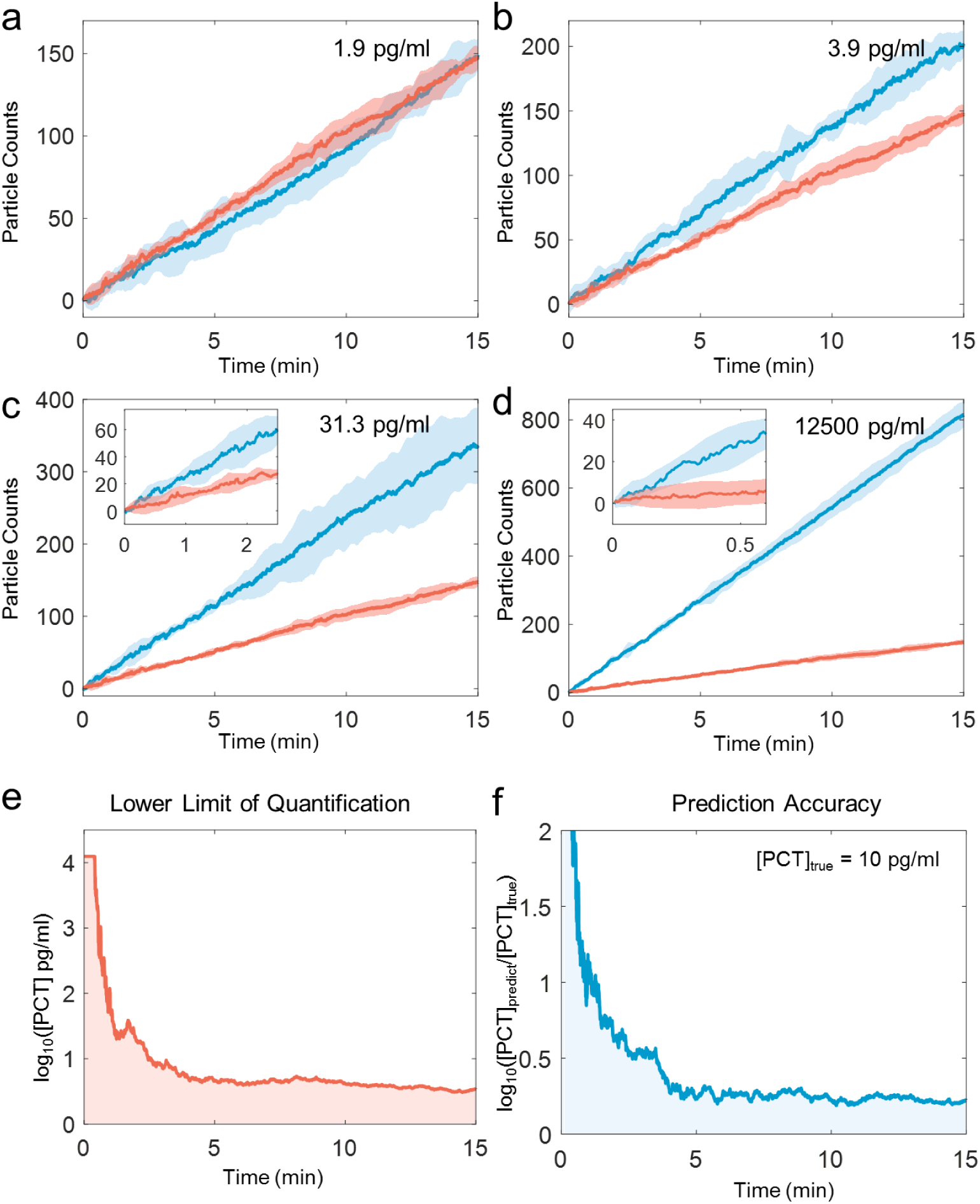
Particle counts vs. counting time at different PCT concentrations. TD-immunoassay measurements of PCT at concentration of 1.9 pg/ml **(a)**, 3.9 **(b)**, 31.3 **(c)**, 12500 **(d)** pg/ml, where the thick blue and red lines are the mean values of three replicates for PCT and blank measurements, respectively, and the light blue and light red regions mark the 95% prediction interval. Insets in (c) and (d) are zoom-in plots for time intervals of 2.5 min at 31.3 pg/mL PCT and 0.5 min at 12500 pg/mL. **(e).** Time dependence of lower limit of quantification, and **(f)** Time dependence of prediction accuracy (b) for PCT TD-immunoassay measurements.

### Comparison of TD-Immunoassay with conventional ELISA method using PCT spiked serum samples

To compare the present TD-Immunoassay method with the conventional ELISA method, PCT spiked serum samples with concentrations of 20, 200, and 2000 pg/mL were measured by the both methods, with each measurement repeated three times. The TD-immunoassay results were obtained with 5 min GNP counting time using the PCT standard curve (Fig. S5). As shown in Table 1, both the TD-Immunoassay and conventional ELISA methods can accurately measure the PCT concentrations between 200 and 2000 pg/mL. However, the conventional ELISA does not have sufficient limit of detection to measure samples with PCT concentrations lower than 200 pg/mL. In contrast, the present method can accurately measure PCT in serum with concentrations as low as 20 pg/mL.

**Table 1.** PCT spiked serum calculated respectively by the standard curve of TD-immunoassay counting and conventional ELISA.

| PCT<br>Concentration<br>(pg/mL) | TD-immunoassay*<br>(pg/mL)<br>(*This work*) | Recovery<br>(%) | Replicates | Conventional<br>ELISA (pg/mL) | Recovery<br>(% ) | Replicates |
| --- | --- | --- | --- | --- | --- | --- |
| 20 | 21.4±0.4 | 107±2.1 | 3 | Not<br>quantifiable | Not<br>quantifiable | 3 |
| 200 | 215.7±30 | 95.5±15 | 3 | 197.63±7.8 | 98.5±3.9 | 3 |
| 2000 | 2010±284.8 | 100.5±14.2 | 3 | 1923.24±114.4 | 96.2±5.7 | 3 |

\*The results of TD-immunoassay are the mean and standard deviation calculated based on the standard curve of TD-immunoassay measurements of PCT at 5 min (supplementary information). Each measurement is replicated three times. Recovery (%) is the measured PCT concentration over the actual PCT concentration, which describes the accuracy of each technique.

### Comparison of TD-Immunoassay with different digital immunoassays

Table 2 compares the performance of different digital immunoassays for PCT detection in terms of sample volume, incubation time, total detection time, limit of detection, limit of quantitation, and dynamic range. The TD-immunoassay platform presented here can detect PCT in 100 μL of serum samples with a detection limit of 2.8 pg/mL and a limit of quantitation of 4 pg/mL for a total of 25 min, and dynamic range of ~ 4 logs. This performance is excellent comparted other sensitive PCT detection technologies listed in the table. The most significant improvement of TD- immunoassay is the reduction of the total assay time from hours to ~ 25 min. This is particularly important for acute diseases, such as sepsis.

## Discussion

TD-immunoassay is based on counting of GNPs binding to a sensor surface. The statistical error of counting is given by 1/√*N*, where N is the GNP count, so the detection precision increases with N. The higher concentration of a biomarker, the faster is the binding rate of GNPs, which means shorter detection time to reach a desired precision. This provides a possibility to detect biomarker by tracking the time required to reach a required precision, rather than by counting GNPs over a fixed time interval regardless the biomarker concentration. Considering that patients with positive sepsis tend to have higher biomarker concentrations than normal people, this time resolved detection approach provides fast diagnosis of the patients’ conditions. In addition to shorten the detection time, time-resolved tracking of single GNPs also helps expand the dynamic range. This is because for highly concentrated biomarker, one can choose to shorten the detection time to avoid saturation associated with too many GNPs binding to the sensor surface.

## Conclusions

We have developed a time-resolved digital immunoassay by combining plasmonic imaging and automated counting of single gold nanoparticles. The high contrast plasmonic imaging allows accurate tracking of each nanoparticle. This capability, together with real-time imaging, enables resolving multiple nanoparticle binding to an area within the diffraction limit, and leads to highly precise detection of single binding events with a wide dynamic range. We validated the principle and demonstrated the performance of the immunoassay using PCT in controlled buffer and in sera with a limit of detection of 2.76 pg/mL, limit of quantification of 4 pg/mL, dynamic range of 4-1.25×10^4^ pg/mL, and detection time of less than 25 mins for low concentration samples (a few pg/mL). Our data also shows that the real-time counting can significantly shorten the detection time for high concentration samples (relevant to sepsis patients) and increase the detection dynamic range. Further studies underway include testing of clinical samples. We anticipate that this time-resolved digital immunoassay is particularly useful for diagnosing and tracking progression of acute diseases (e.g., sepsis and cardiovascular diseases), where rapid and precise biomarker quantification is needed.

## Supporting information

## Acknowledgement

The authors are grateful of support from NIH (1R01GM124335) and NSFC (#21773117) (YW).

