## Supplementary material for "Time-resolved digital immunoassay for rapid and sensitive quantitation of procalcitonin with plasmonic imaging"

### Supporting Information

**S1. Protocol of TD-immunoassay.** Figure S1 shows the protocol of the immunoassay reaction used in the present work for PCT detection. The capture antibody was pre-coated on a sensor surface (gold –coated glass slide) using the steps shown in Fig. S1a. The capture antibody-coated sensor was exposed to 100 µL standard or sample and incubated for 10 min, followed by washing with PBST buffer (Fig. S21b). 100 µL biotinylated PCT detection antibody was then added and incubated for 10 min. After washing 100 µL streptavidin-coated GNP solution ($9.41\times10^8$ GNPs per well) was added (~530 streptavidin molecules are covalently coated on each GNP), during which plasmonic imaging was performed for 5 min.

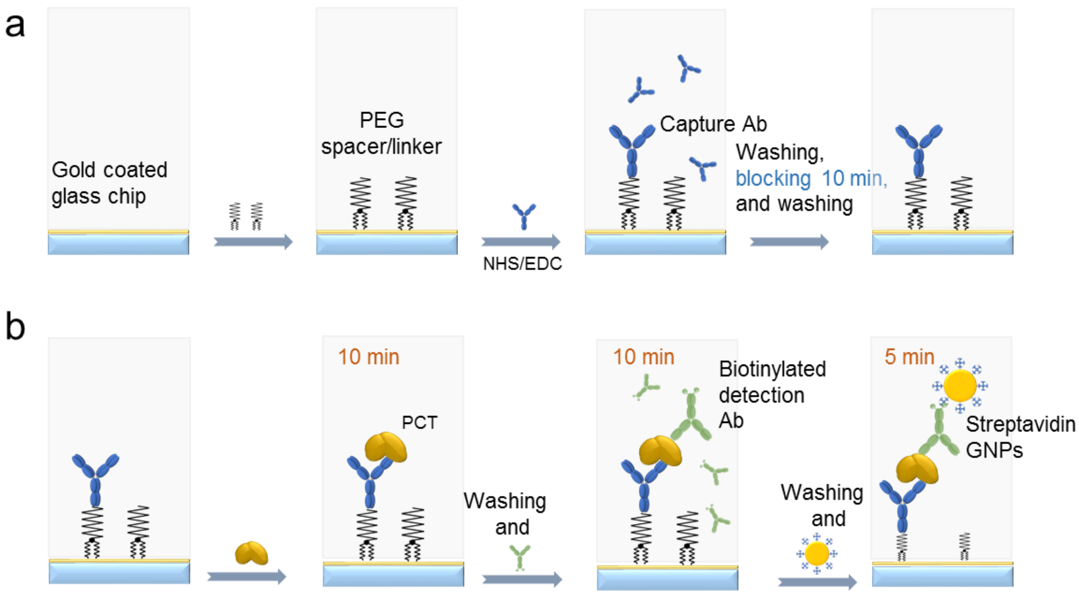


**Figure S1. Protocol of TD-immunoassay. (a)** Preparation of capture-antibody coated sensor surface for PCT detection. **(b)** Steps for the PCT assay with TD-immunoassay.

**S2. Time-resolved tracking of single nanoparticles with plasmonic imaging.** Manual counting of each nanoparticle bound on the gold surface over time is impossible because each test includes 30,000 to 130,000 frames of plasmonic images, and each frame contains a large number of nanoparticles with both binding and unbinding events taking place at the same time. An imaging-processing algorithm was developed to automatically count the binding and unbinding events using the steps shown in Fig. S2. Step 1: Background noise was removed by first applying the K-space filtering process (due to special pattern of GNP plasmonic image [1]), then averaging consecutive blocks and subtracting the previous from the current one to generate differential image sequences. Shot noise was reduced by maximizing the number of photons in the differential images [2,3]. Step 2: To automatically identify and count each nanoparticle in an image, a template was chosen from a pre-processed image sequence and used to correlate with each frame. To eliminate the influence of light intensity, template matching was achieved by normalizing the template and the images to be processed [4]. The binding and unbinding of nanoparticles on each image frame were determined by finding a local maximum in the correlation image. To avoid drift of the images, the algorithm automatically updated the template after a certain period of time. Occasionally, the plasmonic image of a nanoparticle was detected as fluctuating (binding and unbinding). These rare events (<0.1%) were detected but removed from counting. Step 3: From the template-matched patterns on each image frame, nanoparticles associated with binding and unbinding determined, from which the net number of nanoparticles bound to the surface was counted.

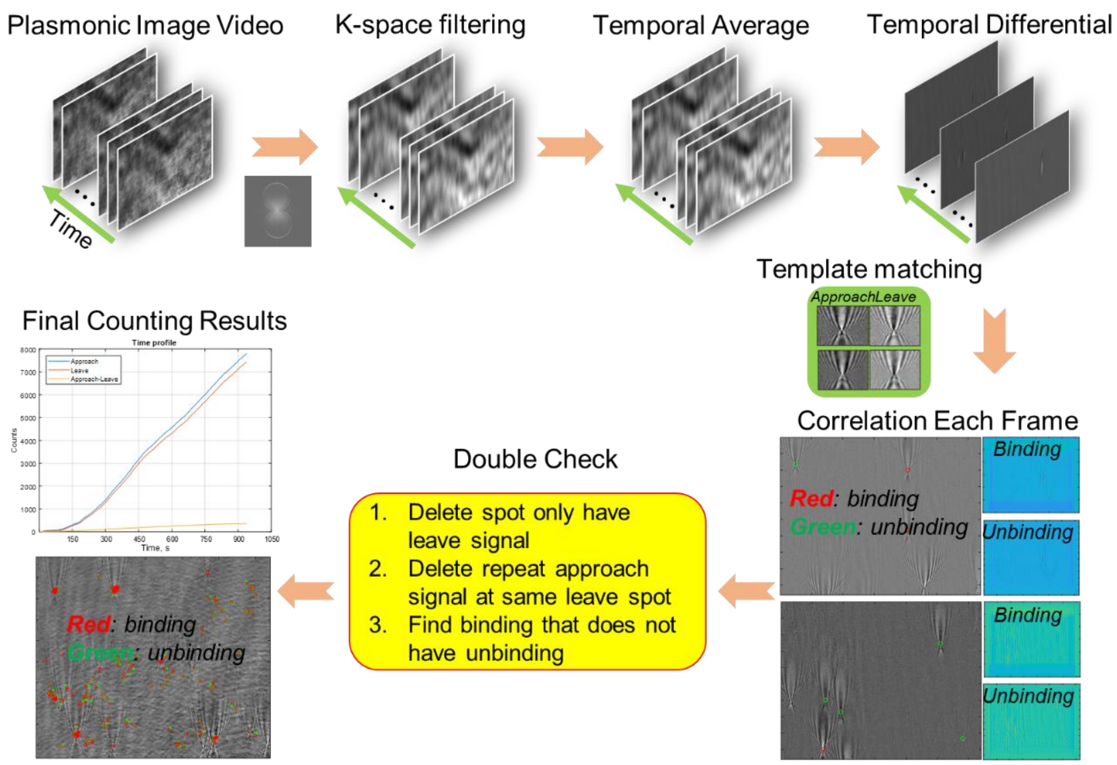


**Figure S2. Automatic particle counting algorithm.** Step 1: Pre-processing plasmonic images using K-space filtering, temporal subtraction and shot noise reduction. Step 2: Identifying nanoparticle with a template-matching algorithm. Step 3: Detect binding and unbinding events. See text for more details.

**S3. TD-immunoassay detection of IgG/anti-IgG binding.** The standard curve of the IgG/anti-IgG binding can be fitted with an empirical equation widely used in ELISA, which is given by

$$y = A_2 + \frac{A_1 - A_2}{1+(\frac{X}{X_0})^p} \qquad (2)$$

with r-square of 0.9986 (Fig. 2e).

**S4. Evaluation of time dependence of TD-immunoassay**

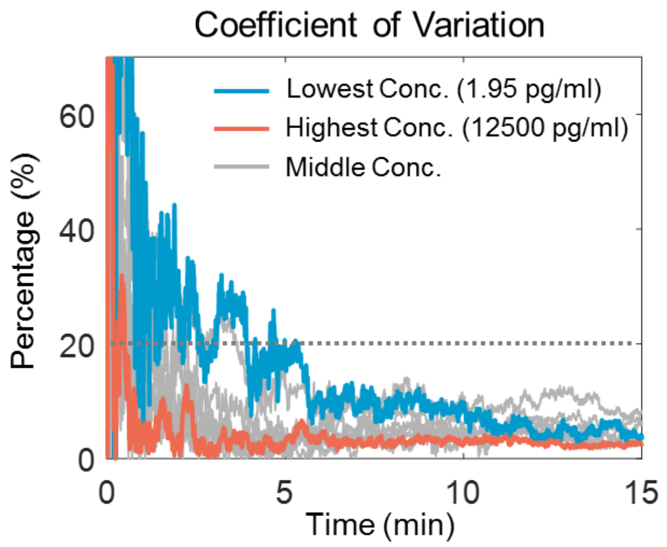


**Figure S3. Evaluation of time dependence of TD-immunoassay.** Time dependence of coefficient of variation for PCT time in the present TD-immunoassay measurements, where the dotted horizontal line indicates 20% coefficient of variation.

**S5. Standard curve of conventional ELISA for PCT detection**

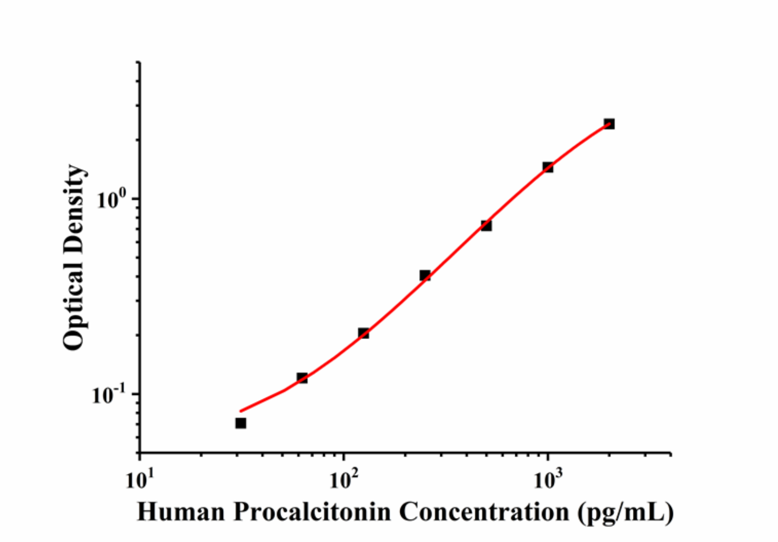


**Figure S4. Standard curve of PCT detection measured by conventional ELISA**. Log-log plot of signal output (r-square = 0,9992 showing a response of 2 logs, range from 31.3 pg/mL to 2000 pg/mL following the instructions of the ELISA kit.

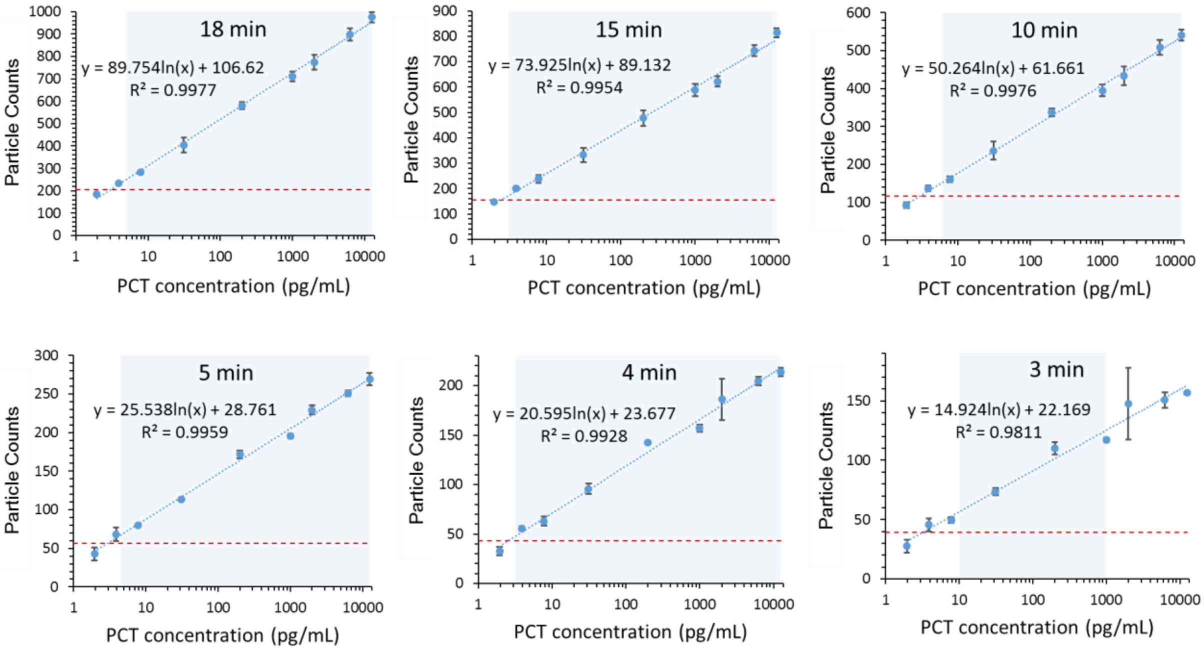

**Figure S5. Standard curve of PCT detection at different time points,** where the horizontal dashed lines show the limit of detection, shaded areas represent the dynamic range, and the error bars are the standard deviation over triplicate test. Because the PCT levels of sepsis patients are usually lower than 10000 pg/mL, PCT concentrations higher than 12500 pg/mL were not tested.

### S6. Dynamic range of TD-immunoassay

TD-ELISA features differential imaging with real-time information, and it quantifies the GNP number change per unit time. The actual dynamic range of TD-immunoassay can be several orders greater than what was demonstrated in the present work. For an imaging frame rate of 26.6 fps, the temporal resolution is 37.6 ms, within which up to $5\times10^3$ GNPs can be resolved if assuming optical diffraction limit (size of diffraction limit used?) for each particle spot. This corresponds to ~$10^5$ GNPs/sec as the upper limit. The lower limit is the detection of a single GNP. For a time interval of 10 min, the lower limit (a single GNP detected in the 10 minutes period) in the detection of GNP binding rate is $1.67\text{x}10^{-3}$ GNP/sec. The full dynamic range of detection is thus about $6\times10^7$. This wide dynamic range covers the requirement of most immunoassay applications. The above discussion assumes 10 min time interval for GNP counting, which can be tuned based on the need of detection time and dynamic range. Because TD-immunoassay also involves incubation of biomarkers with capture antibody, which can also be adjusted based on the need of detection limit, detection time and dynamic range.

### S7. **Statistical Analysis**

We use linear regression model for PCT counting error prediction expressed by

$$N = \beta_0 + \beta_1 \log(c) + \epsilon, \qquad (1)$$

where $N$ is particle counts, $c$ is the PCT concentration, $\beta_0$ and $\beta_1$ are constants and $\epsilon$ describes the error. Given the measurement error $\epsilon$ is mostly affected by the counting error, which depends on particle counts $N$, given by $\epsilon \sim N(0, \sigma^2 Y)$, we transform the heteroscedastic model into a homoscedastic model for the linear regression analysis:

$$\sqrt{N} = \sqrt{\beta_0 + \beta_1 \log(c)} + \epsilon_0,\ \ \epsilon_0 \sim N(0, \sigma^2), \qquad (2)$$

We use 95% prediction interval to evaluate the prediction accuracy from the counting numbers.
